# Hamilton’s rule, the evolution of behavior rules and the wizardry of control theory

**DOI:** 10.1101/2022.06.14.496167

**Authors:** Laurent Lehmann

## Abstract

This paper formalizes selection on a quantitative trait affecting the evolution of behavior (or development) rules through which individuals act and react with their surroundings. Combining Hamilton’s marginal rule for selection on scalar traits and concepts from optimal control theory, a necessary first-order condition for the evolutionary stability of the trait in a group-structured population is derived. The model, which is of intermediate level of complexity, fills a gap between the formalization of selection on evolving traits that are directly conceived as actions (no phenotypic plasticity) and selection on evolving traits that are conceived as strategies or function valued actions (complete phenotypic plasticity). By conceptualizing individuals as open deterministic dynamical systems expressing incomplete phenotypic plasticity, the model captures selection on a large class of phenotypic expression mechanisms, including developmental pathways and learning under life-history trade-offs. As an illustration of the results, a first-order condition for the evolutionary stability of behavior response rules from the social evolution literature is re-derived, strengthened, and generalized. All results of the paper also generalize directly to selection on multidimensional quantitative traits affecting behavior rule evolution, thereby covering neural and gene network evolution.

## 1 Introduction

This paper is about formalizing selection on quantitative traits and exploring mixing two pieces of magic. On one side, there is the magic of Hamilton’s marginal rule, which describes the direction of selection on a quantitative trait in a group (or family) structured population and provides a necessary first-order condition for evolutionary stability (Rousset, 2004; Van Cleve, 2015 for reviews). The magic here, actually mathematics, is that the problem of solving the possibly gigantic system of equations describing the distribution of genetics states among locally interacting individuals is reduced to the much simpler task of computing the probability that two individuals from the same group share a common ancestor under a neutral evolutionary process (all individuals bear the same trait). On the other side, we have the magic of the control theory approach to the calculus of variations, which provides necessary first-order conditions for optimization problems involving objective functions that depend on whole trajectories of dynamical systems (Liberzon, 2011; Weber, 2011 for a reviews), e.g., life-time reproductive sucess depends on development and resource allocation scheduling. The manifold magic here is that the problem of dynamic optimization and solving multidimensional partial differential equations is broken down to the much simpler task of static optimization and computing lower dimensional dynamical systems. In an evolutionary biology context, such partial differential equations capture selection on reactions norms or behavorial strategies, and the analysis can often be reduced to solving ordinary differential equations under a neutral process only.

Both these approaches have opened the door to many applications in evolutionary biology as is illustrated by the vast literatures on social evolution and life-history theories (e.g. Stearns, 1992; Frank, 1998). Wizards have also blended Hamilton’s rule and control theory. In particular, Day and Taylor (1997, 1998, 2000) and Wild (2011) investigated selection on so-called open loop traits under limited genetic mixing. Open loop traits describe phenotypic plastic expression as a function of time (or some other exogenous variable), for instance from birth to death, and represent the standard formalization of reaction norm evolution in life-history theory for panmictic populations. Here, control theory has actually long been applied under the heading of Prontryagin’s maximum principle (e.g., León, 1976; Iwasa and Roughgarden, 1984; Perrin, 1992; Irie and Iwasa, 2005). But plastic phenotypes evolve to vary not only as a function of time or exogenous variables, but also as a function of any fitness relevant state variables determining the physiology, morphology or behavior of an individual, which may themselves depend on phenotypic expression. Such so-called closed loop traits can be thought of as a contingency plans or strategies, since they specify a conditional trait expression rule according to fitness relevant conditions and close the output-input feedback loop between phenotypic expression and individual states. For closed loop traits too, mixtures between Hamilton’s rule and control theory have been explored (Avila et al., 2021), and the evolution of closed loop traits is often studied using dynamic programming (e.g., Houston and McNamara, 1999; Mangel et al., 1988; Ewald et al., 2007; Nakamura and Ohtsuki, 2016).

In the bulk of these formalizations, however, open and closed loop traits display a form of complete plasticity, in the sense that given values of the independent variable(s), time and/or state(s), the phenotype is quantitative and can evolve freely (subject to physiological constraints). As such, the genetically evolving trait is of large dimensions if the independent variables can take many values and evolving traits are typically of infinite dimensions when the range of the dependent variables is continuous. Yet, one can conceive situations of incomplete phenotypic plasticity, where an individual is conceptualized as an open system acting and reacting with its surroundings, but the number of genetic evolving traits affecting phenotypic scheduling is small. Examples actually abound in the literature and include models for the evolution of reactive strategies, behavior response rules, learning rules, or neural and gene networks with a few number of nodes (e.g., Ezoe and Iwasa, 1997; McNamara et al., 1999; Akçay and Van Cleve, 2009; Killingback and Doebeli, 2002; Taylor and Day, 2004; McNamara et al., 2004; André and Day, 2007; Wakano and Miura, 2014; Dridi and Lehmann, 2015; Kobayashi et al., 2019; Alon, 2020). In each of these cases, there is a dynamical system underlying phenotypic scheduling, which, in turn, affects survival and reproduction, and this dynamical system is affected by one or a few number of genetically evolving traits. To a first approximation, most analytical models for reaction norm evolution when interactions occur between individuals are actually of this type. The goal of this paper is to show how standard concepts from control theory, originally devised for situations of complete plasticity where the magic is most effective, are actually also usefull to analyze the evolution of behavioral interactions in group-structured populations under incomplete plasticity. While this topic was partly explored in Avila et al. (2021, their “Constant control result”), some of their results are here generalized. Moreover, life-history scenarios where the analysis turns out be completely analytically tractable are fullly worked out. Finally, a more detailed formalization of an individual interacting with its surroundings is provided. This allows to make explicit connections with the extensive literature on incomplete plasticity and social evolution. The rest of this paper is organized as follows. (i) Hamilton’s marginal rule for selection on scalar traits is recalled and a formalisation of an individual as an open system is provided. (ii) Control theory concepts are used in Hamilton’s marginal rule to derive necessary first-order conditions for the evolutionary stability of a scalar trait affecting any feature of dynamic phenotypic expression of interacting individuals in a life-history evolution contexts. (iii) The generic first-order condition for selection on a trait affecting a behavior response rule in group-structured population (Akçay and Van Cleve, 2012, eq. 5, Akçay and Van Cleve, 2014, eq. 8) is re-derived, strengthened, and generalized. (iv) Limitations and generalities of the model are discussed.

## 2 Model

### 2.1 Main assumptions

#### 2.1.1 The evolutionary setting and Hamilton’s marginal rule

Consider a population of homogeneous individuals (no class structure) that can be subdivided into a large number of groups of fixed size *n* and were the population is censused at discrete demographic time steps, i.e., the standard social evolution model setting as reviewed in Michod (1982) and Rousset (2004). Interactions may occur among individuals within groups and/or between groups before reproduction. These interaction are assumed to be affected by an evolving quantitive genetic trait *u*, which either belongs to the whole set R of real numbers or a subset 𝒰 = [*u*_min_, *u*_max_] thereoff (i.e., *u* ∈ 𝒰 ⊂ ℝ, for instance when *u* is a probability, then *u*_min_ = 0 and *u*_max_ = 1). The interactions are also assumed to be such that a focal (or representative) individual from the population has its survival and reproduction affected by individuals in up to three distinct roles *sensu* Grafen (2006, p. 544) in relation to the focal. Namely, the focal individual itself with trait *u*_*•*_, its group members with average trait *u*_*○*_, and individuals at large from the population with average trait *u*. Individuals in each role are thus different types of actor on the focal individual’s fitness *w*(*u*_*•*_, *u*_*○*_, *u*), which gives its expected number of successful offspring produced over one demographic time step (including self through survival) when expressing trait value *u*_*•*_, when group neighbours express average trait value *u*_*•*_, and individuals from the population at large express average trait value *u* (formally, *w* : *𝒰*^3^ *→* ℝ_+_, which is assumed differentiable and demographic consistency due to density-dependent regulation requires that when the population is phenotypically monomorphic for trait value *u*, fitness is equal to one: *w*(*u, u, u*) = 1 for all *u ∈ 𝒰*).

Let us further denote by *r*(*u*) the neutral relatedness between two randomly sampled individuals from the same group when the population is phenotypically monomorphic for trait value *u*. Thus, *r*(*u*) is the probability that in a neutral process (where all individuals are alike with trait *u*) two homologous genes of these individuals coalesce in a common ancestor (e.g., Michod and Hamilton, 1980; Frank, 1998; Roze and Rousset, 2003; Rousset, 2004; Lehmann and Rousset, 2014; Van Cleve, 2015). A necessary first-order condition for trait value *u*^*∗*^ *∈* [*u*_min_, *u*_max_] to be uninvadable (all mutant deviations go extinct) is that

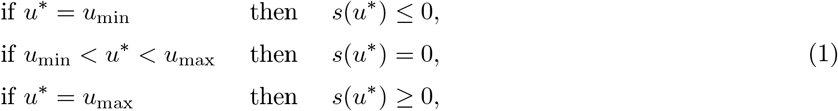

where

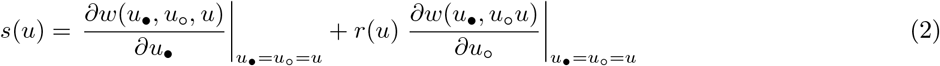

is the inclusive fitness effect from changing trait value in a population monomorphic for *u*. This is the sum of the marginal effect *w*(*u*_*•*_, *u*_*○*_*u*)*/∂u*_*○*_of the focal individual varying (infinitesimally) its trait expression level on its own fitness plus the marginal effect *∂w*(*u*_*•*_, *u*_*○*_*u*)*/∂u*_*○*_ of an average group neighbour on the focal’s fitness weighted by neutral relatedness. This latter effect can also be read in an actor-centered way as the sum of the effect on the fitness of any single neighbour of the focal individual varying its trait expression (Mullon et al., 2016, eq. 12).

This first-order condition for evolutionary stability (eqs. 1–2) is a central result of social evolution theory that has been derived in many different ways, some more precise and detailed than others, from the population genetics of allele frequency change and fixation probabilities, to the Price equation and evolutionary invasion analysis, and to quantitative genetics (see, e.g., Taylor and Frank, 1996; Frank, 1998; Rousset and Billiard, 2000; Roze and Rousset, 2003; Rousset, 2004; Taylor et al., 2007; Akçay and Van Cleve, 2012; Wakano et al., 2012; Lehmann and Rousset, 2014; Mullon et al., 2016; Van Cleve, 2015; Mullon and Lehmann, 2019; Van Cleve, 2020; Avila et al., 2021). All these derivations fully agree with each other and this stubbornly robust result will be taken as the evolutionary backbone and starting point of the present analysis. Eqs. (1)–(2) also imply that an increase in trait value brought by mutation is favored, to the first-order in a resident population at *u*, when *s*(*u*) *>* 0, i.e., Hamilton’s marginal rule holds and a trait value *u* satisfying *s*(*u*) = 0 will be called a singular trait value. Nothing in Hamilton’s marginal rule implies that the evolving trait is a particular behavioral action and this is true for Hamilton’s (1964) initial formulation of the inclusive fitness effect. It could be any quantitative genetic trait affecting more or less remotely morphological, physiological, or behavioral phenotypes potentially affecting interacting individuals under the model’s demographic assumptions. To bring this upfront, we need to first detail phenotypic expression and behavioral interactions and then describe how such interactions affect fitness.

#### 2.1.2 The individual-surroundings interface

Each individual is conceptualized as being an open system exchanging energy, matter, and/or information with its local surroundings possibly at each time point *t ∈* [0, *T*] of the span *T* of behavioral interaction occurring within one demographic time step. This behavioral time span *T* can be thought of as the time available for interactions between individuals. If behavior occurs on a fast time scale relative to reproduction one can take *T → ∞*, i.e., a very large number of interactions occur before reproduction and the separation of times scales between demography and behavior is infinite (a frequently endorsed assumption). An individual as an open system implies that its behavior can be predicted from the knowledge of a set of internal states and a set of external inputs. This perspective is enshrined in animal behavior theory (McFarland and Sibly, 1975; McFarland and Houston, 1981; Enquist and Ghirlanda, 2005) and follows more generally from control theory (or system’s theory e.g., Arbib, 1987; Sontag, 1998; Athans and Falb, 2007; Weber, 2011). As such, the focal individual (section 2.1.1) can be characterized by three phenotypic attributes at each time point *t ∈* [0, *T*] of the behavioral interactions. First, an action *a*_*•*_(*t*) ∈ *𝒜* expressed at time *t*, which is the fundamental phenotypic unit by which the individual interacts with its surroundings, e.g., a motor pattern, a signal, a transfer of resources to another individual, and where *𝒜* denotes the set of actions. Second, an internal state *x*_*•*_(*t*) ∈ *χ*, which is a description of an individual’s internal phenotypic features such as those pertaining to physiology, morphology, or neurology e.g., fat reserve, brain and gonad size, memory and beliefs about locations of food items or family members, and where χ is the set of (internal) states. Third, an external input *ϕ*_*•*_(*t*) ∈ **Φ**, which is any more or less noisy private or public signal about other’s actions or any other cue or material received from the surroundings, e.g., assimilated food, radiant energy, a signal, a perception of the actions of others, where **Φ** denotes the set of inputs. While this state-space representation of behavior is in principle general, even universal (Haykin, 1999), we here make two specific sets of assumptions to obtain a tractable model. First, for simplicity of presentation of the main concepts, we start by assuming that actions and states are one-dimensional real-valued (i.e., *a*_*•*_(*t*) *∈* ℝ and *x*_*•*_(*t*) *∈* ℝ, and the relevant multidimensional case will be discussed in section 4). Second, we assume a deterministic model and that the behavior of the focal at time *t ∈* [0, *T*] can be described by the following system of ordinary differential equations (ODE’s):

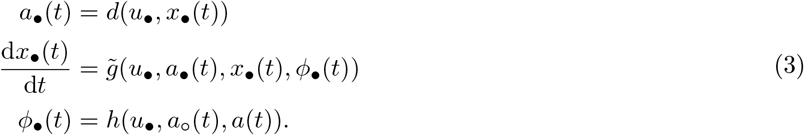

The first line says that the action of the focal individual depends on a decision rule mapping its internal state to action and that this rule is parameterized by the individual’s quantitative trait *𝓊*_*•*_ (formally *d* : *𝒰 ×*ℝ *→* ℝ). The second line says that the change in the state of the focal at time *t* depends on its action *a*_*•*_(*t*) and state *x*_*•*_(*t*), as well as on the input *ϕ*_*•*_(*t*) received from the environment. The mapping 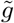 relating these quantities to behavior also depends on the evolving trait (formally 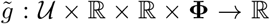). Finally, the last line says that the input *u*_*•*_(*t*) to the focal is a mapping depending on the average action *a*_*○*_ (*t*) of group neighbour, the average action *a*(*t*) in the population at large, and the focal’s genetically determined trait (formally *h* : *𝒰 ×*ℝ^2^ *→* **Φ**). Here and throughout all functions are assumed differentiable.

Eq. (3) specifies how the focal acts and reacts with its surroundings and thus defines its behavior rule *sensu* Lehmann et al. (2015), namely the collection of functions *d*, 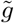, and *h* together with initial conditions for eq. (3). In order to determine the resulting trajectory of actions, we also need to specify the action and state dynamics of group members and individuals at large from the population. Since we assume that individuals are homogeneous and thus differ only by way of expressing different genetic traits, their behavior rules can be taken to be characterized by the same functions *d*, 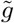 and *h*. Thus, the action of an average group neighbour and average individual from the population are, respectively, given by *a*_*○*_ (*t*) = *d*(*x*_*○*_ (*t*), *u*_*○*_) and *a*(*t*) = *d*(*x*(*t*), *u*). From this and eq. (3), we see that the action and change in state of the focal individual is determined once we know the collections

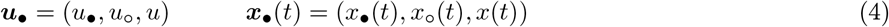

of traits and states, respectively, of all possible actors on the focal’s state dynamics. Thus, it is useful to introduce the function *g* : *𝒰*^3^ × ℝ^3^ *→* ℝ defined as

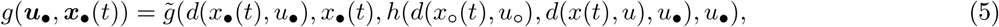

which simply concatenates the different dependencies given in eq. (3). This brings upfront the dependency of the focal’s state dynamics on the collection of traits ***u***_*•*_ and states ***x***_*•*_(*t*) of each actor on the focal’s behavior rule. Thereby, we can write the focal state dynamics as

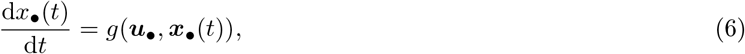

and since all individuals are homogeneous we can express the rate of change of the state of an average neighbour of the focal and an average individual simply by permuting the arguments of the function *g* to obtain

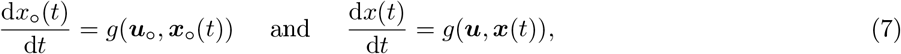

where the vectors

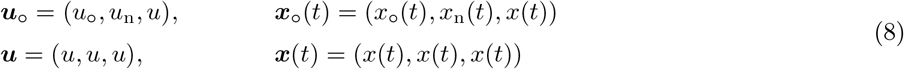

collect the (average) traits and states of actors on the state variables of an average neighbour of the focal individual (first line), and on an average individual in the population (second line), respectively (here and throughout all vectors are defined by default as being column vectors). These actors are thus second-order level actors on the focal recipient since they affect the state variables of actors affecting the focal’s state dynamics. Note that the subscripts of the trait vectors (***u***_*•*_, ***u***_*○*_ and ***u***) and state vectors (***x***_*•*_(*t*), ***x***_*○*_ (*t*) and ***x***(*t*)) emphasise the individual (actor) from who’s perspective the second-order actors’ control and variables are collected. Accordingly, since groups are of size *n*, the vectors in eq. (8) contain the elements

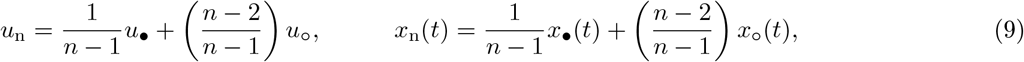

which are, for an average neighbour of the focal, the control and state expressions of average neighbours viewed as actors on the focal individual.

Equations (3)-(7) together fully determine how an individuals acts and reacts with its surroundings. While tracking the dynamics of three state variables (eqs. 6–7) may appear complicated, it is much simpler than tracking the state of all individuals in a group separately plus those in the population, which would require as many equations as there are individuals in the group plus at least the average in the population, and this is the approach often taken in the game theory (e.g., Basar and Olsder, 1999; Weber, 2011) or the evolutionary literatures (e.g., Day and Taylor, 2000). The useful reduction in complexity embodied in eqs. (6)–(7) obtains because using only the average phenotypes and states variable of the different actors on the focal approximates possibly more complicated relationships between genetic states and phenotypic feature to the first-order. Average phenotypes are sufficient to evaluate the selection gradient *s*(*u*), since, to the first-order, expectations of functions depending on phenotypes (or functions thereof) can be replaced by functions of expected phenotypes (or functions thereof) owing to Taylor expansion about mean values [see Rousset (2004, p. 95) for general social evolution considerations, Avila et al. (2021) for application to state space models, and Lynch and Walsh (2018, Appendix 1) for Taylor expansions of functions of random variable in a general context].

#### 2.1.3 Fitness

Having specified how individuals interact, let us now specify how the focal’s fitness *w*(*u*_*•*_, *u*_*○*_, *u*) is determined as a result of these interactions and consider two types of fitness functions. First, the discounted fitness

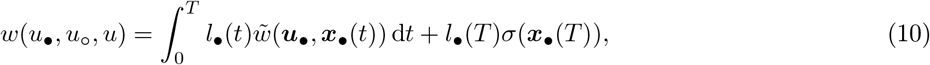

where *l*_*•*_(*t*) is the probability that the focal individual survives until time *t*, which satisfies

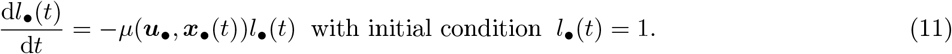

Here, *μ*(***u***_*•*_, ***x***_*•*_(*t*)) is the instantaneous death rate of the focal, which depends on traits and states (since death eventually occurs, we have *l*_*•*_(*t*) *→* 0 as *t → ∞*) and 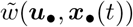 is the rate of increase of fitness due to effective reproduction at time *t* (or the contribution to effective reproduction of the states at time *t*). The term *l*_*•*_(*T*)*σ*(***x***_*•*_(*T*)) is the so-called scrap value, which gives the contribution to fitness from effective reproduction *σ*(***x***_*•*_(*T*)) at the final time *t* = *T* (formally 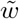 : 𝒰^3^ × ℝ^3^ → ℝ_+_ and *σ* : ℝ^3^ → ℝ_+_). Fitness is actually mediated by actions, since this is the medium by which individuals interact with their surroundings and so the vital rates, 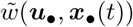 and *μ*(***u***_*•*_, ***x***_*•*_(*t*)), should all depend on actions. But since actions in turn depend on states (recall eq. 3), this justifies to write the vital rates directly as function of the states so that fitness is entirelly determined once we have the solution to eqs. (6)–(7) given initial conditions. Thus, writing fitness in terms of states is a reduced-form expression that implicitly yet fully takes into account the full behavioral model (eq. (3)), which is useful to keep in mind for biological applications.

The discounted fitness eq. (10) covers at least three classes of demographic situations. (a) Interaction between familly members, like between siblings, in a family structured populations (e.g, Michod, 1982; Roze and Rousset, 2004), where individuals can die during the ontogenic period. Depending on the model formulation 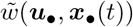 may be zero and all fitness contribution is captured by the scrap value. (b) Interactions among group members in a geographically structured population. With limited dispersal, the fitness component 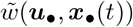 can depends in a non-linear way on various vital rates of group members and individuals from the population at large (e.g, Roze and Rousset, 2003, eq. 35, Akçay and Van Cleve, 2012, eq. A12, Van Cleve, 2015, eq. 38, Mullon et al., 2016, eq. box 1a), which themselves may depend on integrals depending on the traits of the individuals in interaction. Such situations are taken into account in the above formalization either by (i) defining state variables whose integrated values represent the integral, and are covered by the scrap value *σ*(***x***_*•*_(*T*)) in eq. (12) or (ii) by noting that to the first-order, functions of integrals can be replaced by integrals of first-order Taylor series of fitness and hence the 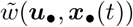 fitness component in eq. (12) may be evaluated as a first-order Taylor expansion of fitness in its vital rates (e.g., Van Cleve, 2015, eq. 39, Mullon et al., 2016, eq. A60-A61). (c) Finally, eq. (10) covers a case that is not fully apparent under the main assumptions (section 2.1.1) yet is useful to mention. This is the case of the standard life-history evolution context in a panmictic age-structured population where the vital rates 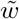 and *μ* depend only on the state of the focal *x*_*•*_(*t*) and its own trait *u*_*•*_ (e.g., Stearns, 1992; Perrin, 1992) and possibly the trait *u* of the population at large to capture density-dependent regulation [or on *u, x*_*○*_ (*t*) to capture interactions among individuals of the same age, e.g., Day and Taylor, 1997]. In this cases, relatedness is zero relatedness (*r*(*u*) = 0), 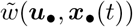 is simply the effective fecundity of the focal individual at age *t*, and 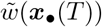 is usually considered nil.

Second, we consider the average fitness

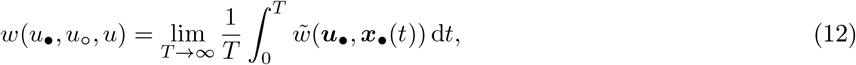

which is the average of the instantaneous contribution 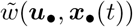 to fitness. Since *T → ∞*, the interactions during behavioral time are assumed to occur on a faster time scale than reproduction. Another way of interpreting this assumption biologically is to say that the behavioral dynamics is able to reach an equilibrium before reproduction occurs. The average fitness eq. (10) can be applied to demographic situations (a)-(b) described above, but does not cover the standard life-history evolution context [demographic situation (c) above], since survival is implicitly assumed to be equal to one.

### 2.2 Selection on behavior rules

#### 2.2.1 The Hamiltonian and the co-state variables

Having specified the assumptions about behavioral interactions (eq. 3 and eq. 6) and fitness (eq. 10 and eq. 12), we now turn to work out a meaningfull representations of the inclusive fitness effect *s*(*u*) using concepts from optimal control theory (Bryson and Ho, 1975; Athans and Falb, 2007; Sydsaeter et al., 2008; Weber, 2011). For this, let us introduce the so-called Hamiltonian function defined here as

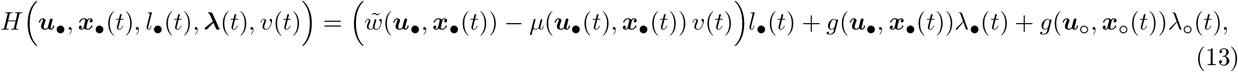

where ***λ***(*t*) = (*λ*_*•*_(*t*), *λ*_*○*_ (*t*)). Extending a classical interpretation (Dorfman, 1969, p. 822), the Hamiltonian (formally the function *H* : *U*^3^ × ℝ^3^ × ℝ × ℝ^2^ × ℝ *→* ℝ) can be regarded as the contribution to the focal individual’s expected fitness of the expression of own and others’ current actions, holding the expression of all future actions constant and thus at their behavioral level determined by the resident trait *u*. The Hamiltonian is thus the sum of the focal individual’s expected current fitness 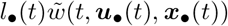 plus the changes of its survival [*−μ*(***u***_*•*_(*t*), ***x***_*•*_(*t*))*l*_*•*_(*t*)] and fitness relevant states [*g*(***u***_*•*_, ***x***_*•*_(*t*)) and *g*(***u***_*○*_, ***x***_*○*_ (*t*))], resulting from current actions, were each such change is weighted by its marginal effect on the focal’s expected remaining fitness. These marginal effects on fitness, *v*(*t*), *λ*_*•*_(*t*) and *λ*_*○*_ (*t*), are collectively referred to as the costates. The first, *v*(*t*), is actually Fisher’s (1930) reproductive value (change in remaining fitness at *t* resulting from change in survival until *t*) while ***λ***(*t*) = (*λ*_*•*_(*t*), *λ*_*○*_ (*t*)) will be called the shadow values of the actions (change in expected remaining fitness stemming from change in states at *t*). All costates are evaluated in a monomorphic resident population where all individual bear trait value *u* (see Box 1 and Avila et al., 2021 for more formalities and interpretations about how the costates connect to reproductive value).

##### Box 1.

**Shadow values and Fisher reproductive value as marginal values**

In order to understand the biologically useful interpretation of the costates, consider the discounted fitness case (eq. 10) and let us introduce the *future value reproductive value* defined as

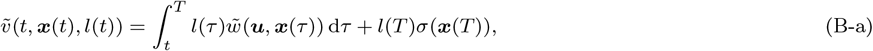

which depends on the whole set of values (or trajectory) of states 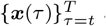 and survival 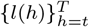 from time *t* to *T* evaluated at *u*. Thus, 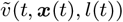 is the expected remaining fitness from time *t* onwards of an individual in a monomorphic population at *u* when the initial conditions for its state is ***x***(*t*) and survival is *l*(*t*). These are the two arguments that, besides time, are emphasized in the notation 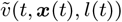 because changes in these initial values will change the future reproductive value by changing the whole trajectories of states and survival starting at *t*, which are themselves functions of the initial states. By contrast, Fisher’s reproductive value, noted *v*(*t*, ***x***(*t*)), is the expected remaining effective fitness from time *t* onwards conditional on the individual surviving until time *t* (Fisher, 1930; Goodman, 1982) and it is therefore given as the *current value reproductive value*:

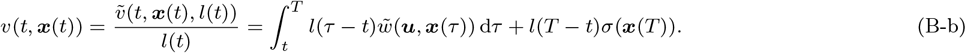

The second equality was reached by noting that 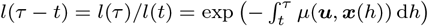is the probability of survival from time *t* to time *τ ≥ t* conditional on surviving until *t*. This shows that *v*(*t*, ***x***(*t*)), similarly to 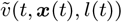, depends on the state trajectory 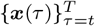 from time *t* to *T*. Yet by contrast to 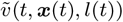, the reproductive value *v*(*t*, ***x***(*t*)) depends on the survival trajectory 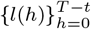 starting at *l*(0) = 1. Since the initial value *l*(0) = 1 is fixed, it does not make biological sense to vary it and this explains why the argument of *v*(*t*, ***x***(*t*)), besides time, only involves ***x***(*t*) whose variation will indeed affect *v*(*t*, ***x***(*t*)). Hence, it follows from 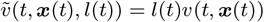 that

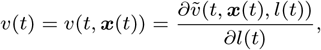

which is the usual partial derivative of 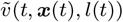 with respect to *l*(*t*). In other words, Fisher’s reproductive value is the marginal change of the future value reproductive value with respect to change in survival (see also León, 1976; Perrin, 1992). Such interpretation of the costates variables as partial derivatives, namely, as marginal changes in future value reproductive value when the initial state variable values are changed holds generally (Sydsaeter et al., 2008), since one also has

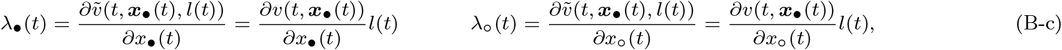

where all partial derivatives are evaluated at *u* and ***x***_*•*_(*t*) = ***x***(*t*) = (*x*(*t*), *x*(*t*), *x*(*t*)). The second equality follows from using eq. (B-b) under the form 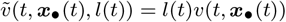 [see Avila et al., 2021, Appendix B.1 for a proof that, in an evolutionary context, costates are generally the partial derivatives of 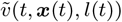 with respect to state change regardless of the mode of trait control]. Finally, it can be useful to note that demographic consistency entails that

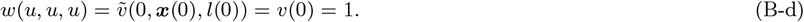

Further, under the infinite horizon case (*T → ∞*) for constant reproduction and mortality, say 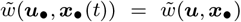 and *μ*(***u***_*•*_, ***x***_*•*_(*t*)) = *μ*(***u, x***_*•*_(*t*)), one has 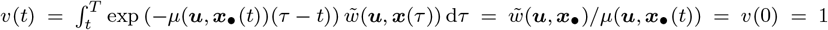 for all *t*.

The Hamiltonian thus captures the fundamental trade-off between current and future fitness consequence of expressing current actions and delineates all pathways that can affect this trade-off by bringing upfront the fitness value of changing states. The Hamiltonian also allows to generate the rate of changes of the costates, which are obtained by differentiation: d*v*(*t*)*/* d*t* = *−∂H/∂l*_*•*_(*t*), d*λ*_*•*_(*t*)*/*d*t* = *−∂H/∂x*_*•*_(*t*), and d*λ*_*○*_ (*t*)*/* d*t* = *−∂H/∂x*_*○*_ (*t*), where all partial derivatives, here and throughout, are evaluated at ***u***_*•*_ = ***u*** = (*u, u, u*) and ***x***_*•*_(*t*) = ***x***(*t*) = (*x*(*t*), *x*(*t*), *x*(*t*)). From this and using eq. (9) in eq. (13), the costates are fully determined by the following system of ODE’s:

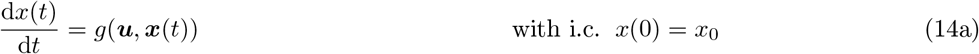

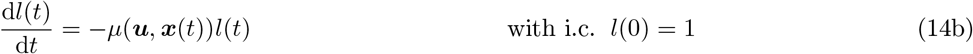

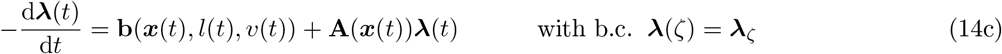

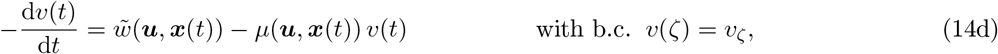

where

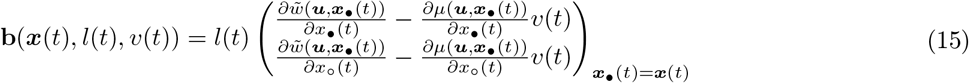

is a vector of marginal changes in fitness stemming from changes in state and

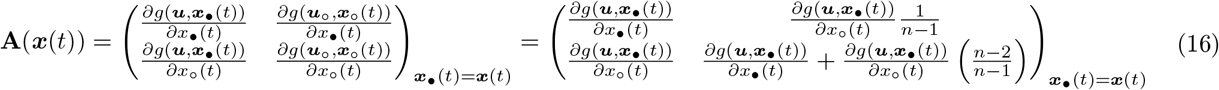

is a matrix of changes in state dynamics. The second representation of **A**(***x***(*t*)) follows from using eq. (9), which allows to express **A**(***x***(*t*)) only in terms of the changes in state dynamics of the focal individual and both representations of **A**(***x***(*t*)) in eq. (16) can be usefull. In system (14), “i.c” stands for initial condition, i.e., the value of the states at time *t* = 0 and “b.c.” stands for boundary condition, i.e., the value of the costates at some arbitrary time *t* = *ζ*. This is due to the fact that, depending on the situation, the boundary conditions of the costates are initial or final values (more on this below). While eq. (14a) will generally be non-linear, eq. (14c) is a linear non-homogeneous ODE with varying coefficients and the representation of its solution is discussed in Appendix A.1.

The ODE system (14) is centerpiece to the analyzes of selection and given initial condition for the states, boundary conditions for the costates, and trait value *u*, its solution is unique. So system (14) can be seen to be parameterized by *u* and while the initial condition of the states, *x*_0_ is a parameter as well, the boundary conditions of the costates are determined by the specificities of the situation at hand. No general recipe for determining the boundary conditions exists (e.g., Aseev and Kryazhimskiy, 2008; Sydsaeter et al., 2008), yet for the finite horizon case, the boundary conditions are straighforwardly obtained from their definitions as marginal effects of changes in states on remaining fitness evaluated in a monomorphic population at *u*: *v*(*T*) = *σ*(***x***(*T*)) and 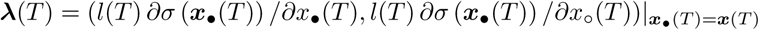 (see Box 1). Difficulties for determining the boundary condition arise for discounted fitness under the infinite horizon case, where it may be felt that the costates go asymptotically to zero but this is not generally the case (Sydsaeter et al., 2008; Aseev and Kryazhimskiy, 2008; Weber, 2011). For instance, for constant survival and reproduction, the reproductive value remains the same throughout lifespan and the reproductive value has the constraint that its initial value is one (see Box 1). In the presence of *dominating discount*, however, where a constant discount rate suppresses the growth rate of the expected current fitness, the growth rate of trajectory *x*(*t*), as well as that of the trajectory of the corresponding regular linearized dynamical system, then the shadow values should go to zero; namely lim_*T→∞*_ ***λ***(*T*) = (0, 0), which fixes their boundary conditions (Aseev and Kryazhimskiy, 2008; Aseev and Veliov, 2019). Since external mortality can usually be taken as a constant, there is alway a background of constant discount to potentially suppress the growth rates. Further, because physiological state variables are bounded and thus cannot grow indefinitely, it is actually plausible that the majority of biological situations of interest do satisfy the dominating discount assumption, at least by assuming high enough external mortality, one may be able to enforce it for a large class of biological scenarios. This is proved in Appendix A.2, where the conditions leading to lim_*T→∞*_ ***λ***(*T*) = (0, 0) are discussed more formally and biologically. Now fully endorsing dominating discount, we are lead to our first result about the representation of the selection gradient proved in Appendix B.2.1

### Result 1

#### Discounted fitness with finite and infinite horizon

*Suppose individual fitness is given by eq. (10) with state dynamics by eqs. (6)-(7), then the inclusive fitness effect for condition* (1) *can be represented as*

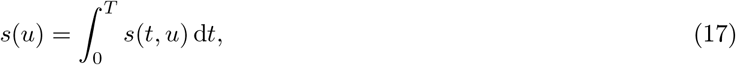

*where*

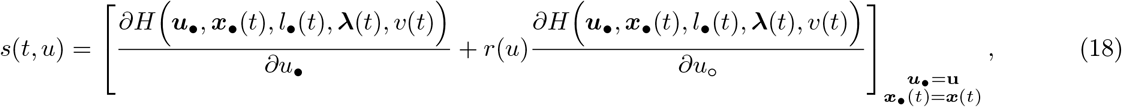

*which, owing to eq*. (13), *can also be written as*

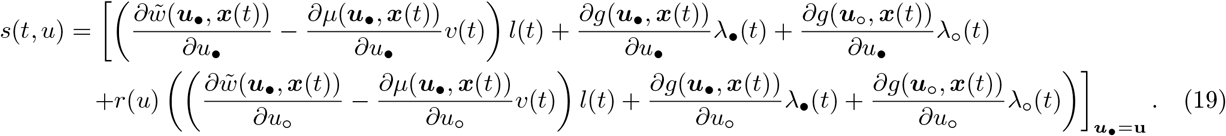

*Therein, the states and costates satisfy eq. (14) with b*.*c. of the costates given by v*(*T*) = *σ*(***x***(*T*)) *and* 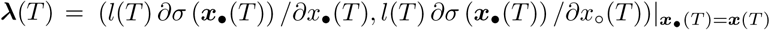 *for a finite horizon. For an infinite horizon, eq. (17) holds with the assumption that* lim_*T→∞*_ ***λ***(*T*) = (0, 0), *i*.*e*., *dominating discount obtains*.

Result 1 generalizes to the infinite horizon case and to an explicit behavioral rule evolution context the Constant control result of Avila et al. (2021, p. 11). From a biological perspective, it says that the selection gradient on the evolving trait is the sum over all times, of the inclusive effect of trait change on expected current and remaining reproduction. The latter effect is mediated by how the change in trait value changes own and others’ state, weighted by the marginal effect of changing these states on expected remaining reproduction. It is useful to note that the selection gradient can also be expressed solely in terms of the (Fisher) reproductive values by factoring out *l*(*t*) from *s*(*t, u*), which yields

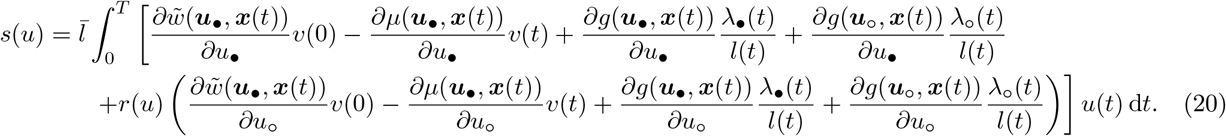

Here, 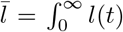 is the average lifespan of an individual and 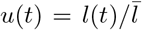 is the probability that a randomly sampled individual from the population is of age *t*. According to eq. (B-c) of Box 1, *λ*_*•*_(*t*)*/l*(*t*) and *λ*_*○*_ (*t*)*/l*(*t*), are the changes in reproductive value stemming from change in own and other’s state at *t*, respectively. Hence, *s*(*u*) consists of two terms connected to reproductive value. First, the two first summands on each line of eq. (20) together give the expectation over all ages in which an individual can reside of the reproductive value weighted changes of vital rates induced by inclusive trait change effect (hence the delineation of the offspring reproductive value *v*(0) = 1, Box 1). This is conceptually equivalent to the standard representations of the selection gradient in heterogeneous populations (e.g., Lion, 2018, eq. 21 for panmictic populations and Priklopil and Lehmann, 2020, eq. 2 for metapopulations). Second, the last two summands on each line together give the expectation over all ages in which an individual can reside of the changes in reproductive value resulting from state changes induced by inclusive effects of trait change. This second term arises owing to the dynamic constraints on state and is thus bound to the first term through the common currency reproductive value. In the absence of such constraints, eq. (20) is fully consistent with the classical selection gradient on a mutation that affects survival and/or fecundity in a panmictic populations and obtained from the geometric growth rate of the mutation (Ronce and Promislow, 2010, eqs. 3.1–3.3 by noting that their scalling of reproductive values, i.e., their eq. A-11, implies that for a continuous time model 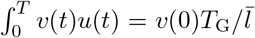, where *T*_G_ is the mean generation time, a scaling factor allowing to covert the first-order perturbation of the basic reproductive number into that of the geometric growth rate). Finally, eq. (20) is also consistent, as it should, with the representation of selection on life-history traits to any order and given by the Hamilton-Bellman-Jacobi equation expressed in terms of (Fisher) reproductive value only (León, 1976, eq. 36).

From, a computational perspective, result 2 shows that the main technical difficulty to evaluate condition (1) is to solve the ODE system (14) and integrate *s*(*t, u*) over time. This is generally a much simpler task than attempting to solve the problem by “brute force”, i.e. solving eqs. (6)–(7), substituting into fitness (eq. (10) or eq. (12)) and then differentiating. Hence, there are gains to use the control theory concepts for both obtaining a biological interpretation of the selection gradient on a trait under incomplete plasticitiy and for the concrete computation of that gradient. Depending on the model, however it may remain a challenge to solve the ODE system (14) and we now describe a case where this can be markedly simplified. This is the case where the biological situation dictates that the initial state satisfies the resident equilibrium, i.e., 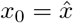 with 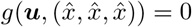 where ***u*** = (*u, u, u*). While this clearly excludes many situations pertaining to life-history evolution, it does include biologically relevant cases. For instance, it obtains when the initial condition of the states are inherited from the previous generation, which occurs in the case where knowledge, resources, or stationary behavior, is transferred between generations and this is typical of models for the evolution of learning or social interactions (e.g., Killingback and Doebeli, 2002; Taylor and Day, 2004; Aoki et al., 2012; Wakano and Miura, 2014; Kobayashi et al., 2019, see also section 3). To analyze this case, let 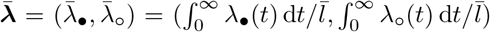 be the average shadow values over lifespan (or average of the change of Fisher reproductive value). This leads us to the following result proved in Appendix B.2.2.

### Result 2

#### Discounted fitness with infinite horizon and initial resident equilibrium state

*Suppose individual fitness is given by eq. (10) with state dynamics by eqs. (6)-(7) under an infinite horizon with dominating discounts, i*.*e*., *T → ∞ with* lim_*T→∞*_ ***λ***(*T*) = (0, 0), *and that the initial state* 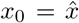 *satisfies* 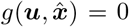 *where* 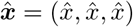. *Then, the inclusive fitness for condition* (1) *can be written as*

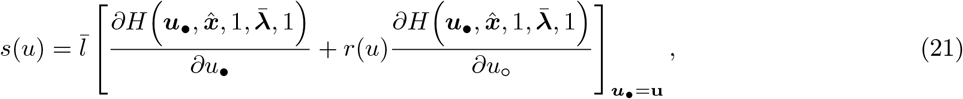

*or more explicitly as*

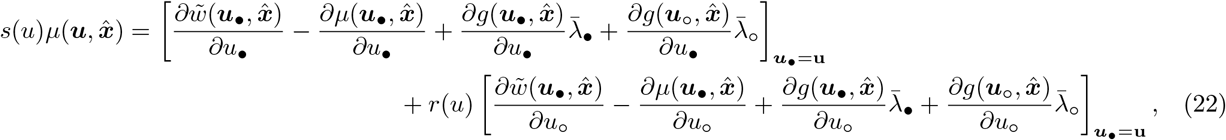

*where* 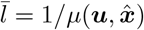 *and* 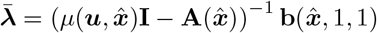 *with* **I** *being the identity matrix*.

From a biological perspective, this result says that the selection gradient on the trait is the sum of the inclusive effects of trait change on the current growth rate 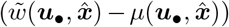 and on reproductive value averaged through lifespan. This latter effect is mediated by how the change in trait value changes own and others’ state expression weighted by the marginal effect of the state changes on average reproductive value. From, a computational perspective, there is no difficulty at all in evaluating *s*(*u*), since it only requires inversion of a 2 by 2 matrix. This result makes analytically tractable behavior rule evolution models when there is a trade-off between effects on fecundity and mortality. Let us finally turn to a closely related result proved in Appendix B.2.3, which is the representation of the selection gradient under average fitness (12).

### Result 3

#### Average fitness

*Suppose individual fitness is given by eq. (12) with state dynamics by eqs. (6)-(7) and that given initial conditions x*(0) = *x*_0_ *and* ***λ***(0) = ***λ***_0_ *at t* = 0, *the dynamics* (14a) *and* (14c) *converge to the (hyperbolically stable) equilibrium* 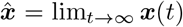 *and* 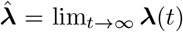. *Then the inclusive fitness effect for condition* (1) *can be written as*

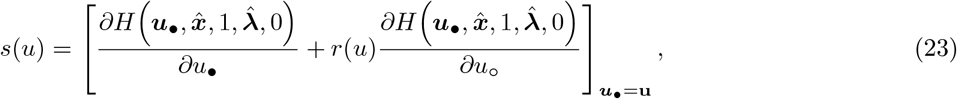

*or more explicitly as*

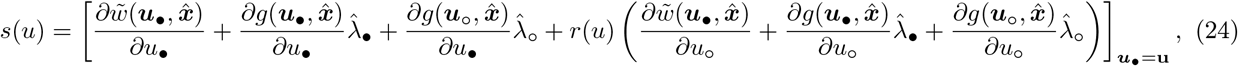

*where* 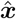 *satisfies* 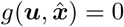 *and the costates* 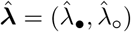 *satisfy* 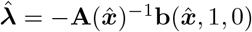.

From a biological perspective, this result says that the selection gradient is the sum of the inclusive effects of trait change on stationary current and future reproduction. The latter effect is mediated by how the change in trait value changes own and others’ future state values weighted by the marginal effect of state change on stationary future reproduction. This is so because the average fitness-(12) entails that only the equilibrium value of the state variable and shadow value will matter for fitness (a consequence of the ergodic theorem). From, a computational perspective, we are in the same straightforward position to evaluate explicitly *s*(*u*) as under result 2.

These results 1-3 can be seen from at least two perspectives. First, they allow to analyze new situations for the evolution of phenotypic scheduling under various social evolution and life-history contexts by providing a recipe to compute the selection gradient on the trait affecting phenotypic dynamics. Second, they provide a biological interpretation of selection on such traits using control theory concepts, which can be used to get insights about the selection pressure in previous evolutionary models under incomplete plasticity. In the next section, we turn to illustrate these points.

## 3 Application: behavioral response models

We now illustrate the results by applying them to the evolution of behavior response rules, in which one or a few scalar traits affect how individuals interact with each other under various type of games (e.g. McNamara et al., 1999; Killingback and Doebeli, 2002; Taylor and Day, 2004; Lehmann and Keller, 2006; André and Day, 2007; Akçay and Van Cleve, 2009; Akçay and Van Cleve, 2012). To make the connection to this literature, we make the main assumptions implied by the model formulation in this literature for the control theory framework. First, states correspond directly to actions; namely the decision rule is the identity function (*a*_*•*_(*t*) = *x*_*•*_(*t*)) and thus does depend on the evolving trait. Further, the input is simply the action of group neighbours (*ϕ*_*•*_(*t*) = *a*_*○*_ (*t*)). Second, the transition rule describing change of action depend only on the actor’s evolving trait; namely *g*(***u***_*•*_, ***x***_*•*_(*t*)) = *g*(*u*_*•*_, ***x***_*•*_(*t*)) and *g*(***u***_*○*_, ***x***_*○*_ (*t*)) = *g*(*u*_*○*_, ***x***_*○*_ (*t*)). Third, reproduction and survival depend directly only on state variables and not on the evolving trait of the actor; namely

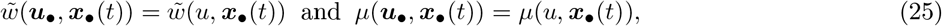

where the dependence on the resident trait *u* must remain owing to density-dependent regulation (i.e, the iron rule that *w*(*u, u, u*) = 1). Together, the second and third assumption imply that

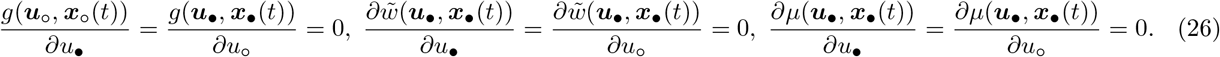

This setting covers more or less implicitly models for the evolution of so-called reactive strategies in the iterated prisonner’s dilemma game or other repeated games (e.g. McNamara et al., 1999; Killingback and Doebeli, 2002; Taylor and Day, 2004; Lehmann and Keller, 2006; André and Day, 2007; Akçay and Van Cleve, 2014), certain models for the evolution of preferences in games (e.g. Heifetz et al., 2007; Akçay and Van Cleve, 2009; Alger and Weibull, 2012; Akçay and Van Cleve, 2012), as well as models for the evolution of learning in games (e.g. Dridi and Akçay, 2018; Leimar, 2021). This is so because all formalize the evolution of behavior rule parameterized by one or a few quantitative genetic traits and it can further be usefull to note that a large class of (stochastic) learning dynamics can actually be described by differential equations fitting into the setting of the behavioral rule model eq. (3) (Fudenberg and Levine, 1998; Dridi and Lehmann, 2014).

### 3.1 Average fitness

Let us consider selection on a trait affecting the evolution of a behavioral response rule following assumptions (26) by using average fitness (eq. 12). This setting should cover the model analyzed in Akçay and Van Cleve (2012) (that generalizes Akçay and Van Cleve, 2009 to interactions between relatives and *n* players) since this model is premised on the facts that (i) individuals interactions occur among groups of fixed size *n* in a geographically structured population; (ii) the dynamical system underlying individual actions has reached stationarity (Fig. 1 and eq. 12 of Akçay and Van Cleve, 2012); (iii) this dynamical system is influenced only by the evolving genetic trait of that individual (eq. 2 of Akçay and Van Cleve, 2009 and implied by the analysis in Akçay and Van Cleve, 2012); and (iv) the fitness/payoff depends only on state variables having reached equilibrium and not directly on the evolving traits (first equation on p. 260 of Akçay and Van Cleve, 2012). This is thus an ideal situation to check the consistency of the optimal control approach to behavior rule evolution, since we should be able to recover the first-order condition for uninvadability of Akçay and Van Cleve (2012, eq. 5).

Under these assumptions, the analysis of the model is covered by result 3. Substituting eq. (26) into eq. (24) shows that the selection gradient reduces to

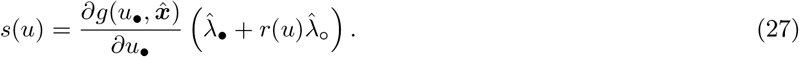

Selection thus depends on how the change in trait of the focal individual affects the change in its action dynamics, 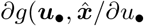, weighted by 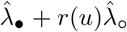, which can be interpreted as the stationary inclusive shadow value of the focal’s actions. This is the sum of the effect 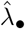 of the focal’s action on its (stationary) fitness and the effect 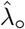 of these actions on the (stationary) fitness of an average neighbour weighted by relatedness *r*(*u*) between the two individuals. A singular trait must satisfy 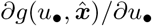 and/or 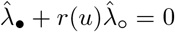. Since the behavior response rule literature usually assumes that the function *g* is monotonic in its first argument, we focus on characterizing the condition 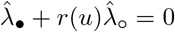.

Under assumptions (25)–(26) and from eq. (14c), the equilibrium shadow values satisfy

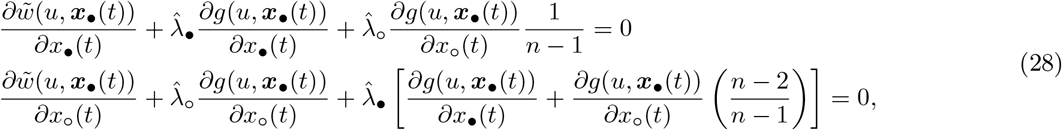

where recall that all derivatives are evaluated at 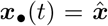. Note that for the behavior response rule setting, the first column of matrix **A**(***x***(*t*)) (recall eq. (16)) describes the change in action of the focal individual when its own action and that of its neighbours are varied [first and and second entry, respectively, given by *∂g*(*u*, ***x***_*•*_(*t*))*/∂x*_*•*_(*t*) and *∂g*(*u*, ***x***_*•*_(*t*))*/∂x*_*○*_ (*t*) in eq. (28)]. The second column of **A**(***x***(*t*)) describes the change of action of a neighbour of the focal when the focal and its neighbours change action. Matrix **A**(***x***(*t*)) can thus be interpreted as describing the action reaction to action between interacting individuals and since the equilibrium shadow values satisfy 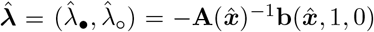, the equilibrium action interdependence is captured by matrix 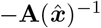. The ratio *ρ*(*u*) = *a*_12_*/a*_11_, where *a*_*ij*_ stands for entry *ij* of matrix 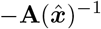, can thus be regarded as a behavioral response coefficient that tells us how much, in equilibrium, the action of a single neighbour of the focal is varied when the focal varies its action relative to the focal’s own induced action variation. Algebraic rearrangements (easily done with a symbolic algebra system) then show that the response coefficient can be expressed as

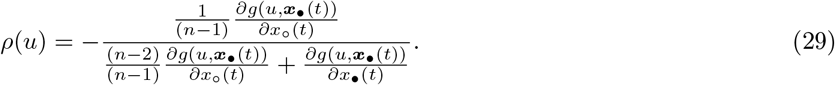

Now solving for 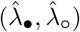 in eq. (28), substituting into eq. (27), and rearranging shows that the condition for *u* to be singular can be written as

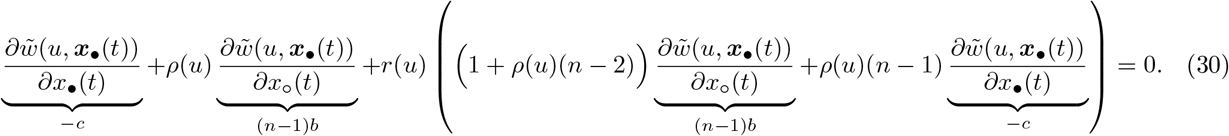

This first-order condition turns out to be conceptually exactly equivalent to eq. (5) of Akçay and Van Cleve (2012) [and eq. (8) of Akçay and Van Cleve, 2014], since we can write 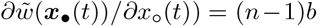, where *b* is conceptually equivalent to the effect of Akçay and Van Cleve (2012) of a single neighbor changing action on the focal’s fitness and the response coefficient (29) is actually exactly eq. (13) of Akçay and Van Cleve (2012). This can be seen by noting that *∂g*(**u, *x***_*•*_(*t*))*/∂x*_*○*_ (*t*) is the effect of the whole set of group neighbours on the action dynamics of the focal individual, while *∂*^2^*x*_*j*_*/*(*∂a*_*j*_*∂a*_*i*_) is the effect of a single neighbour on the action dynamics of the focal in Akçay and Van Cleve (2012) and so *∂g*(**u, *x***_*•*_(*t*))*/∂x*_*○*_ (*t*)*/*(*n −* 1) = *∂*^2^*x*_*j*_*/*(*∂a*_*j*_*∂a*_*i*_) (the special notation *∂*^2^*x*_*j*_*/*(*∂a*_*j*_*∂a*_*i*_) stems from the fact that Akçay and Van Cleve, 2012 endorse a gradient dynamics).

Since the response coefficient can be interpreted as the effect of the focal changing action on the action of a single neighbour (see also eq. 2 of Akçay and Van Cleve, 2012), this paves the way to an intuitive actor-centered interpretation of eq. (30) by considering that 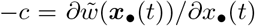 is a fitness cost. Namely, eq. (30) says that the fitness cost to self of varying (infinitesimally) action expression can be recouped by two pathways. First, by the sum of the effect of each neigbour changing actions on the focal fitness weighted by the extent *ρ*(*u*) to which any neighbour’s action is varied when the focal varies its action. Second, by the net indirect effect on the fitness of any related neighbour, which consists of three distinct fitness effects: (i) the immediate effect of the focal varying action on the fitness of each related neighbour [effect of intensity *r*(*u*)(*n −* 1)*b*]; (ii) the correlated effect resulting from the change of action of each neighbour of a focal’s neighbour when the focal varies its action, which induces a change in fitness in any of the focal’s related neighbour [effect of intensity *r*(*u*)*ρ*(*u*)(*n −* 2)(*n −* 1)*b*]; and (iii) the correlated effect of the change of action of each related neighbour of a focal’s neighbour when the focal varies its action, which changes those neighbors fitness [effect of intensity *−r*(*u*)*ρ*(*u*)(*n −* 1)*c*]. As emphasized by Akçay and Van Cleve (2014), this entails an interesting but neglected interaction between relatedness and behavioral responses, owing to the product *ρ*(*u*)*r*(*u*) appearing in eq. (30).

Three other facts about the first-order condition eq. (30) are noteworthy. First, its original derivation by Akçay and Van Cleve (2012) is more direct and simpler. Namely, it consist of letting the state first go to equilibrium and use fitness at that state as 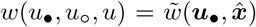 (which can be obtained from eq. (12) by invoking the ergodic theorem). Then, using implicit differentiation of the growth rates of actions at equilibrium with respect to trait values eventually leads to eq. (30). Second, eq. (30) is written at the fitness levels and the units of measurements therein are thus different than those of eq. (5) of Akçay and Van Cleve (2012), since they do not consider effective fecundity 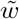, but actual fecundity. As such *r* in their eq. (5) is not relatedness but the scaled relatedness coefficient (e.g., Van Cleve, 2015, eq. 74, Alger et al., 2020, eq. 22) for an iteroparous population. This connects to the point of the last paragraph of section 2.1.3: given some specific life-cycle assumptions express fitness in terms of vital rates and then use the chain rule to express fitness effects in terms of vital rates effect and then one would recover eq. (5) of Akçay and Van Cleve (2012) with the same units of measurement. In other words, eq. (30) and eq. (5) of Akçay and Van Cleve (2012) are scaled differently. Third, while preference evolution was emphasized in the original derivation of condition (30), it can as well be used to study the evolution of reactive strategies (e.g., Killingback and Doebeli, 2002; Taylor and Day, 2004; André and Day, 2007), which is a connection that is worked out in Akçay and Van Cleve (2014) and so will not be repeated here.

More generally, the first-order condition (30) also applies to the evolution of traits affecting learning dynamics. In fact, it applies to any model that can be put under the form of the behavior rule model eq. (3) [along with assumptions (25)–(26)] and so is robust to a multitude of assumptions about behavioral dynamics. Hence, depending on the biological context at hand, evolution my lead to quite different values of the response coefficient *ρ*(*u*) and it is not a conclusion that *ρ*(*u*) always evolves to be positive. For instance, results on preference evolution under incomplete information (Alger et al., 2020) implies that *ρ*(*u*) goes to zero and results under complete information (e.g., Heifetz et al., 2007) imply that it can take a variety of intermediate values depending on the nature of the interactions between individuals and the constraints on preferences. On the other hand, models of reactive strategies suggest that *ρ*(*u*) can sometimes go to one (André and Day, 2007). Hence, the values to which *ρ*(*u*) evolves depends on assumption about phenotypic expression mechanisms as well as the type of interactions individuals face.

### 3.2 Discounted fitness

Let us now consider selection on a trait affecting the evolution of a behavioral response rule under assumptions (26) by using discounted fitness under an infinite time horizon (eq. 10 with *T → ∞*) under the assumptions of Result 2. This entails that the initial condition of the state 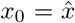 satisfies 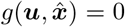, so that the behavioral dynamics is at steady state. Then, the analysis is covered by result 2, and on substituting eqs. (25)–(26) into eq. (22) shows that the the selection gradient reduces to

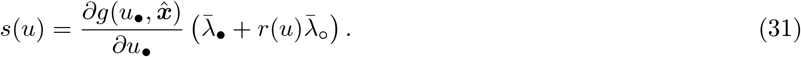

Hence, selection now depends on how the change in trait affect the change in the action dynamics of the focal individual, 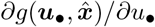 weighted by the inclusive average shadow value of these actions. As for the selection gradient in the previous section (eq. 27), let us focus on the condition where 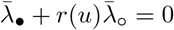. To characterize this in terms of the expressions of the shadow values, note that, as in the previous section, matrix **A**(***x***(*t*)) describes the action reaction to action between interacting individuals. Yet individuals may die as they age and for this model, the average action interdependence is captured by the matrix 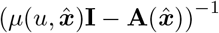 since 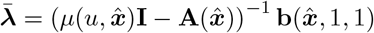. As such the ratio 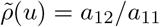, where *a*_*ij*_ now stands for entry *ij* of matrix 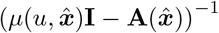, can be regarded as the behavioral response coefficient that tells us how much, on average throughout the whole span of interactions, the action of a single neighbour of the focal is varied when the focal varies its action relative to the focal own induced action variation. Algebraic rearrangements show that the response coefficient is now given by

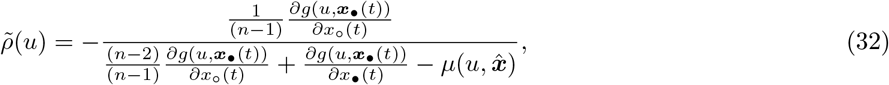

and the difference with eq. (29) is thus only the 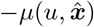 in the denominator. This will act to reduce the value of the response coefficient (owing to the minus sign in front of the ratio). This makes biological sense: the change of action by an individual may not feedback on future fitness through change of action by neighbours if the individual dies.

Using assumptions (25)–(26) in eqs. (15)–(16), computing 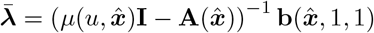, substituting into eq. (31), and rearranging shows that the condition for *u* to be singular can be written as

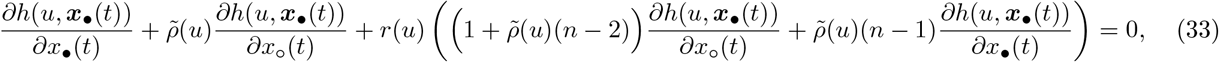

where

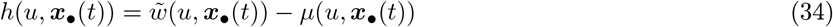

is the stationary growth rate of an individual with trait value *u*. Equation (33) has exactly the same structural form as eq. (30), and so we don’t need to repeat its interpretation, since everything carries over *mutatis mutandis* from eq. (30). The difference, however, is that one can now study behavior response evolution analytically in the presence of survival-fecundity trade-offs and thus embed such evolution in a more explicit life-history context to analyze how behavioral responses depend on demographic variables. We can also conclude that the structural feature of the selection gradient derived by Akçay and Van Cleve (2012) is robust to changes in a number of demographic assumptions. And it is now appropriate to circle back to their assumption of the separation of time scales between behavioral dynamics and demography. This assumption could be one way to justify the assumption that the initial states satisfy the behavioral equilibrium (needed to use eq. 31). In other words, behavior dynamics is so fast that equilibrium is reached before any event, such as death, can occur. Alternatively, individuals could be born in a population that has reached the behavioral equilibrium and inherit that from their parents. This suggests that, as long as there are no interactions between the convergence towards the behavioral equilibrium and the demographic variables themselves, the selection gradient on traits underlying response rules should take the functional form given by the left hand side of eq. (30).

## 4 Discussion

This paper combined Hamilton’s marginal rule for scalar quantitative traits and concepts from optimal control theory to derive the selection gradient on a trait affecting the phenotypic scheduling of an individual interacting with its surroundings. The different first-order conditions (results 1-3) allow to study selection on behavior rules under incomplete plasticity for a reasonably large class of different scenarios under (i) evolution in homogeneous group (or family) structured populations and (ii) life-history evolution under panmixia.

As an application and sanity check of the results, the seminal first-order condition for behavior response rule evolution in group structured populations of Akçay and Van Cleve (2012, eq. 5) was re-derived and generalized to broader behavioral and life-history contexts. This illustrates that the control theory approach provides an overarching framework to formalize selection on quantitative traits affecting phenotypic expression and scheduling that feed back on survival and reproduction. Control theory concepts straddle the layers of plasticity and complexity, since they can be applied to models of complete plasticity ranging from standard open and closed loop traits to models of incomplete plasticity as considered here. And for each case, Hamilton’s rule is seamlessly blended in to consider evolution in spatially structured populations (Avila et al., 2021).

An obvious limitation of the present formalization is that it applies only to homogeneous group-structured populations and it would thus be relevant to think about developing it for heterogeneous group-structure and possibly isolation-by-distance. This is in principle doable since control theory can be applied to age-structured populations that are also class-structured (Avila et al., 2019). Another limitation is that the model is deterministic and thus does not consider stochastic effects on state dynamics. Such effects may not only impact state dynamics direclty but also indirectly through environmental stochasticity, which will impact environmentally dependent state variables. In order to include stochasticity one could replace the system of ordinary differential equations (ODE’s) for the state dynamics by stochastic differential equations (SDE’s), and then attempt to apply results from stochastic control theory (e.g. Basar and Olsder, 1999; Kamien and Schwartz, 2012; Fleming and Soner, 2006) to generalize the first-order conditions to the case where fitness involves expectation of integrals over lifespan. This opens avenues for future research.

For ease of presentation of the concepts, the two central phenotypic attributes of the model, the state *x* and evolving trait *u*, were assumed to be one dimensional. Is anything changed if these variables take values in some finite multidimensional space; namely, if state *x* = (*x*_1_, *x*_2_, …) is a finite vector with *x*_*i*_ being the *i*-th state variable and trait *u* = (*u*_1_, *u*_2_,..) is a finite vector with *u*_*i*_ being the *i*-th quantitative trait? For this case, the selection gradient on trait *u*_*i*_ is still given by the inclusive fitness effect (eq. 2). Further, all other equations of the paper are actually maintained functionally, one just need to carefully read all one-dimensional mappings as multidimensional ones, and any product of scalars as the corresponding dot product between vectors if the scalar has a vectorial counterpart. But no new concept is needed and therefore results 1-3 directly generalize to finite multidimensional traits, states, and actions.

An interesting and perhaps biologically surprising fact that is gained by going from the one-dimensional to the finite multidimensional case, is that this is sufficient to make the model universal in the sense that any behavior rule can in principle be modeled with it. The reason is that a finite multidimensional state dynamics is sufficient to describe the dynamics of a finite recurrent neural network, and the neural network weights can be taken as the evolving quantitative traits. In turn, finite recurrent neural network can implement any type of behavior since they are computationally equivalent to Turing machines (Siegelman and Sontag, 1995; Haykin, 1999). In other words, evolving neural networks allow to consider in principle the evolution of any behavior rule, and here one could let the level of plasticity itself be evolving. This could also open avenues for future research.

It has long been stressed that constructing more explicit mechanistic models for the evolution of phenotypes would be usefull (e.g. McNamara and Houston, 1999; Lotem, 2012; Fawcett et al., 2012; Akçay, 2020). Perhaps one reason this has not been systematically undertaken is the lack of the identification by evolutionary biologists of a coherent framework that allows to analyze adapative models across a wide spectrum of mechanistic formulations. Control theory blended with adaptive dynamics–“evolutionary control theory”–provides just that. My hope is thus that the present formalization and more generally the many spellbinding insights from control theory (e.g., Bryson and Ho, 1975; Arbib, 1987; Sontag, 1998; Haykin, 1999; Dockner et al., 2000; Liberzon, 2011; Athans and Falb, 2007; Sydsaeter et al., 2008; Astrom and Murray, 2008; Weber, 2011) can be used as sources of inspirations to attempt to close the loop between mechanistic and functional aspects of phenotypic expression evolution.

## Acknowledgments

I thank M. Cant for a discussion that crucially stimulated the writing of this paper, P. Avila for comments on the manuscript, and C. Mullon for repeated usefull discussions. I also thank the two reviewers for their very constructive comments.

## Appendix A Cauchy formula

### Appendix A.1 Cauchy formula and Fisher reproductive value

We here give a representation of the solution to the non-homogeneous linear system of ordinary differential equations of the form d***λ***(*t*)*/* d*t* = *−***b**(*t*) *−* **A**(*t*)***λ***(*t*) with b.c. ***λ***(*ζ*) at *t* = *ζ*. This covers the shadow value dynamics, eq. (14c) of the main text, since for that case **b**(*t*) = **b**(***x***(*t*), *l*(*t*), *v*(*t*)) and **A**(*t*) = **A**(***x***(*t*)). According to the Cauchy formula, the following solution then holds for all *t ∈* [0, *T*]:

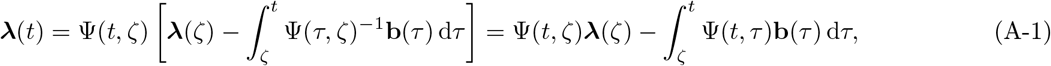

where Ψ(*t, τ*) is the so-called fundamental matrix satisfying

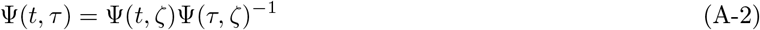

as well as the initial value problem

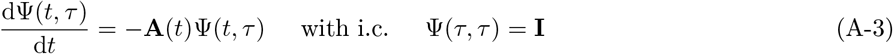

with **I** being the identity matrix (e.g., Athans and Falb, 2007, pp. 127–130, Weber, 2011, pp. 69–72, Aseev and Kryazhimskiy, 2008, eq. 15). The *i*th column of Ψ(*t, τ*) is the solution ***λ***(*t*) of the system d***λ***(*t*)*/* d*t* = *−***A**(*t*)***λ***(*t*) when the initial condition ***λ***(*τ*) is the standard basis with unit value at entry *i*, zero otherwise. For the one dimensional case, the solution is

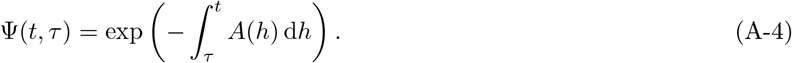

For the multidimensional case, there is no analytic representation for Ψ(*t, τ*) when the system has variable coefficients and various method have been developed to evaluate the fundamental matrix Ψ(*t, τ*) (Weber, 2011, p. 71). For the constant coefficient case, where **A**(*t*) = **A** for all *t*, we get the standard matrix exponential form solution

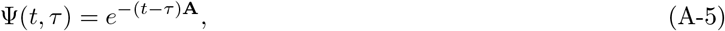

whereby eq. (A-1) reduces to

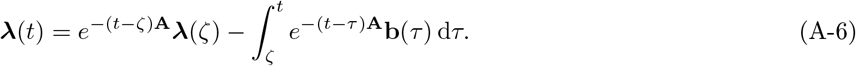

The above results can be used to solve for the reproductive value dynamics 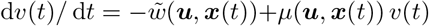 under the finite horizon case (recall eq. 14d), which, in terms of the above notation, is obtained by setting 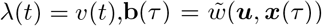, and **A**(*t*) = *−μ*(***u, x***(*t*)). Then taking *ζ* = *T* as the final condition with *v*(*T*) = *σ*(***x***(*T*)) and recalling that 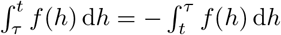, we see that by substituting eq. (A-4) into eq. (A-1), we get

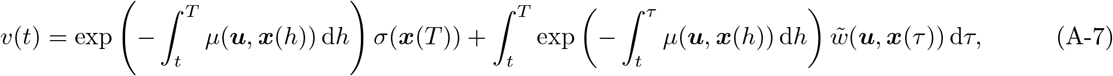

which is exactly the expression for the reproductive value given in Box 1, as it should.

### Appendix A.2 Cauchy formula and dominating discount

We here use the insights and adapt the argument of Aseev and Kryazhimskiy (2008) and Aseev and Veliov (2019) to characterize the condition under which lim_*t→∞*_ ***λ***(*t*) = (0, 0) when fitness takes the infinite horizon discount form 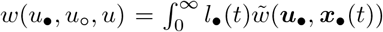 (eq. (10) with *T → ∞*). To do this, let us first write the death rate of the focal individual as *μ*(***u***_*•*_, ***x***_*•*_(*t*)) = *μ*_e_ + *μ*_i_(***u***_*•*_, ***x***_*•*_(*t*)), where *μ*_e_ *∈* ℝ_+_ is the external mortality and *μ*_i_(***u***_*•*_, ***x***_*•*_(*t*)) is the part of the death rate that is endogenously (or internally) determined by the evolving trait and the states.

The term *μ*_i_(***u***_*•*_, ***x***_*•*_(*t*)) could be positive or negative but is assumed to never suppress the external mortality, i.e., *μ*_e_ + *μ*_i_(***u***_*•*_, ***x***_*•*_(*t*)) *>* 0. With this, the discounted fitness can be expressed as

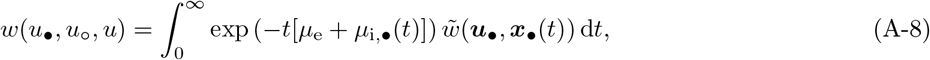

where 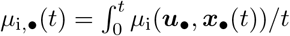 is the average death rate until time *t*.

Now, setting **b**(*τ*) = **b**(***x***(*τ*), *l*(*τ*), *v*(*τ*)) and **A**(*τ*) = **A**(***x***(*τ*)), and taking the boundary condition *ζ* = *∞* and assuming that ***λ***(*∞*) = (0, 0) in the Cauchy formula (A-1), we get 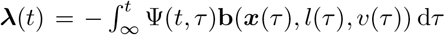. Interchanging the boundaries and using eq. (A-4) at *ζ* = 0, the representation of dynamics becomes the solution of the shadow values

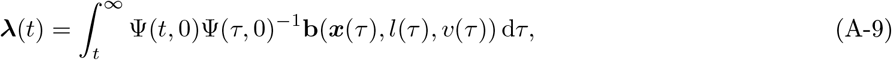

where

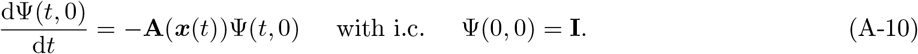

The remaining goal is to determine the conditions under which lim_*t→∞*_ ***λ***(*∞*) = (0, 0) in eq (A-9) such that it is consistent with the assumption ***λ***(*∞*) = (0, 0) that was put into its derivation and so eq (A-9) indeed satisfies eq. (14c) with boundary condition ***λ***(*∞*) = (0, 0). For this, we make the following assumptions.

i. There exist numbers *C*_1_ *≥* 0 and *C*_2_ *≥* 0 such that

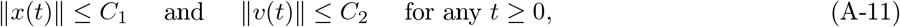

where, here and throughout ∥·∥ denotes an appropriate norm. This assumption says that the value of the state dynamics and the reproductive value are bounded. Bounded reproductive value is biologically realistic and if *x*(*t*) is a morphological or physiological state variable, bounded state variable is also realistic.
ii. There exist numbers *C*_3_ *≥* 0, *C*_4_ *≥* 0, *r*_1_ *≥* 0, and *r*_2_ *≥* 0 such that

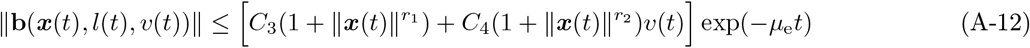

for any admissible ***x***(*t*) and *v*(*t*). The terms 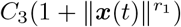 and 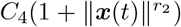 in this condition ensures that 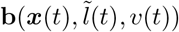, which can be seen as the gradient of expected current conditional individual fitness 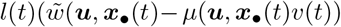 with respect to change of states is not too abrupt (i.e., the slope is not infinite; the first term accounts for the slope of 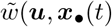, while the second for that of *μ*(***u, x***_*•*_(*t*)). In evolutionary models, this is generally the case when the value of the state variables is bounded away from zero. The multiplier exp(*−μ*_e_*t*) indicates that fitness (A-8) and thus 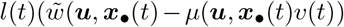 contains the death rate *μ*_e_. Condition (ii) appears applicable to a broad class of evolutionary models and it would be interesting to document a model where it does not hold. Owing to condition (ii), ineq. (A-11) actually reduces to

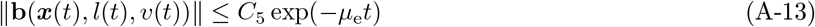

for some constant *C*_5_ *≥* 0.
iii. Let the non-homogeneous linearized dynamical system

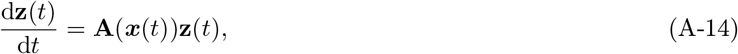

for the state variable be regular. This means that the sum of the two characteristic exponents of the (Lyapunov) spectrum of system (A-14) is equal to the average, over time, of the traces of the sequence of **A**(***x***(*t*)) matrices (Aseev and Kryazhimskiy, 2008, p. 524). If system (A-14) has a constant matrix, like under the assumptions of result 2, then it is necessarily regular. More generally, if the process ***x***(*t*) is ergodic so that the sequence of matrices **A**(***x***(*t*)) is ergodic, then system (A-14) is regular (Pikovsky and Politi, 2016, p. 22). Then for any *ϵ >* 0, as small as needed, the following inequality holds

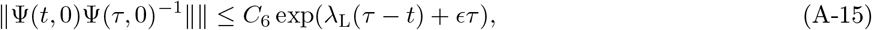

where *λ*_L_ is the largest characteristic exponent of system (A-14) and *C*_6_ *≥* 0 is a constant depending only on *ϵ* (Aseev and Kryazhimskiy, 2008, eq. 28, Aseev and Veliov, 2019, eq. 72).

With these assumptions in hand, let us recall that owing to (a) the relationship between norms and integrals (Athans and Falb, 2007, eq. 3-111) and (b) the relationship between norms of matrices and vectors (Cohen, 2003, p. 16), we have from eq. (A-9) that

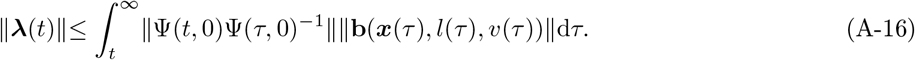

Then, using eqs. (A-12)–(A-15) into eq. (A-17), we get

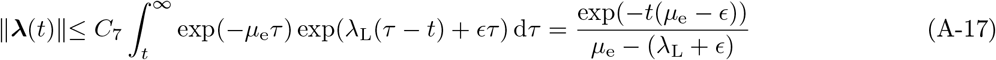

for some constant *C*_7_ *≥* 0 and where the equality holds if *μ*_e_ *> λ*_L_ + *ϵ*. Since *ϵ* can be choosen as small as required, the condition

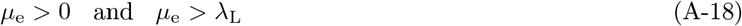

ensures the validity of lim_*t*→∞_∥**λ**(t)∥= 0 and thus lim_*t→∞*_ ***λ***(*t*) = (0, 0).

In summary, if state variables are bounded, fitness components do not change too abruptly, and the external mortality rates suppresses the growth rate of the regular linearized dynamical system of the states variables, then the shadow values go to zero, i.e., lim_*t→∞*_ ***λ***(*t*) = (0, 0). It is plausible that a majority of evolutionary models where state variables represent internal factors fall into this category, hence the claim of the main text. For the cases falling outside this category, it is relevant to know that Aseev and Kryazhimskiy (2008) and Aseev and Veliov (2019) provide conditions on the discount rate under which the shadow value still takes the usefull representation (A-17) even if lim_*t→∞*_ ***λ***(*t*) ≠ (0, 0).

## Appendix B Selection gradient

### Appendix B.1 Generic expression for the selection gradient

We here give the generic representation of the inclusive fitness effect *s*(*u*) when fitness takes the form of eq. (10) or eq. (12), and the state dynamics satisfy eqs. (6)-(7). The representation of *s*(*u*) for these cases is implicitly contained in the calculations of Avila et al. (2021, Appendix B.2.1), who derived an expression for the selection gradient in the broader context where the evolving trait itself is dynamic and of infinite dimensions, thus covering generally both open-loop and closed-loop traits as well. But since this more broader context involves more complicated mathematics and notations, the represention of *s*(*u*) is here (re)-derived for clarity and completeness.

To that end, note first that the survival probability *l*_*•*_(*t*) can itself be thought of as a state variable and eq. (10) of the main text can then be written as

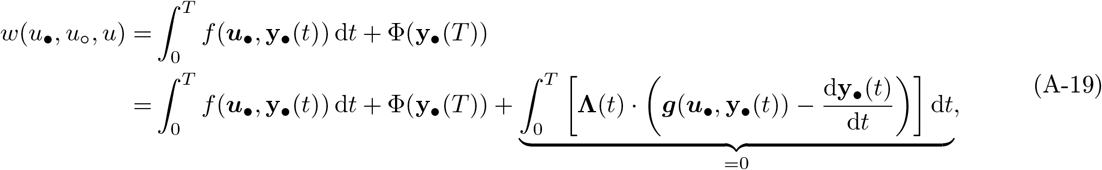

where 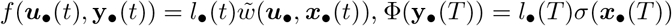, the vector **y**_*•*_(*t*) = (***x***_*•*_(*t*), *l*_*•*_(*t*)) = (*x*_*•*_(*t*), *x*_*○*_ (*t*), *l*_*•*_(*t*)) collects all the state variables, the vector **Λ**(*t*) = (*λ*_*•*_(*t*), *λ*_*○*_ (*t*), *v*(*t*)) collects all costates, and the vector d**y**_*•*_(*t*)*/* d*t* = ***g***(***u***_*•*_, **y**_*•*_(*t*)) = (*g*(***u***_*•*_, ***x***_*•*_(*t*)), *g*(***u***_*○*_, ***x***_*○*_ (*t*)), *−μ*(***u***_*•*_(*t*), ***x***_*•*_(*t*))*l*_*•*_(*t*)) collects all changes in states. The symbol *·* denotes the dot product between two vectors and integrating the last term in eq. (A-19) by parts yields

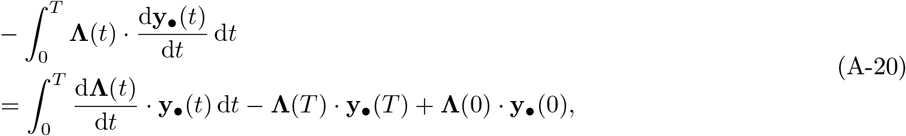

and hence eq. (A-19) becomes

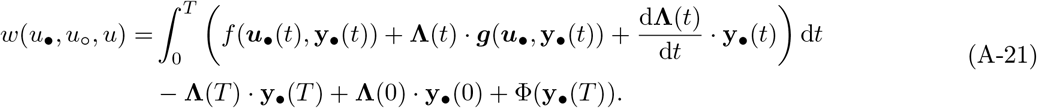

Using the Hamiltonian

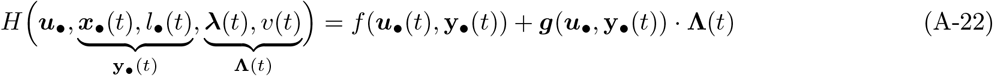

(eq. (13) of the main text), and

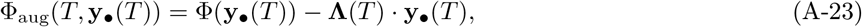

allows to write eq. (A-21) as

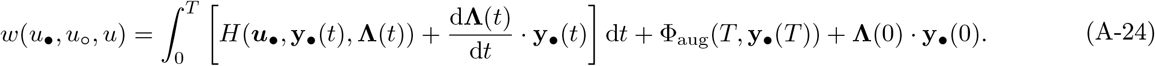

Taking the derivative of fitness (A-24) with respect to the focal’s trait *u*_*•*_ yields in force of the chain rule and Leibniz’s formula (Sydsaeter et al., 2008, p. 160) that

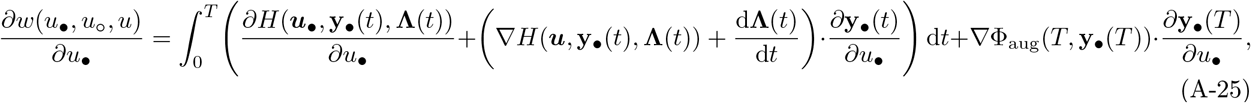

where the gradient operator *∇* is with respect to variable **y**_*•*_(*t*), e.g., *∇H* = (*∂H/∂x*_*•*_(*t*), *∂H/∂x*_*○*_ (*t*), *∂H/∂l*_*•*_(*t*)), *∂***y**_*•*_(*T*)*/∂u*_*•*_ = (*∂x*_*•*_(*T*)*/∂u*_*•*_, *∂x*_*○*_ (*T*)*/∂u*_*•*_, *∂l*_*•*_(*T*)*/∂u*_*•*_), and all derivatives, here and throughout, are evaluated at ***u***_*•*_ = ***u*** = (*u, u, u*) and **y**_*•*_(*t*) = (***x***(*t*), *l*(*t*)) = ((*x*(*t*), *x*(*t*), *x*(*t*)), *l*(*t*)). Likewise, we have

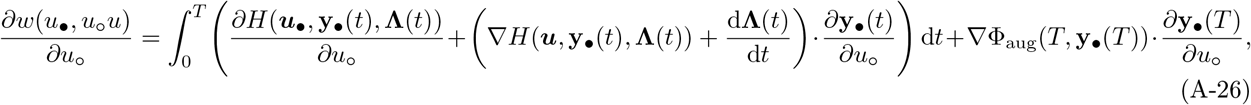

where *∂***y**_*•*_(*T*)*/∂u*_*○*_ = (*∂x*_*•*_(*T*)*/∂u*_*○*_, *∂x*_*○*_ (*T*)*/∂u*_*○*_, *∂l*_*•*_(*T*)*/∂u*_*○*_).

Note that the derivatives of the term **Λ**(0) *·* **y**_*•*_(0) in eqs. (A-25)–(A-26) have vanished because the initial state variables *x*_*•*_(0) = *x*_*○*_ (0) = *x*(0) = *x*_init_ do not depend on the evolving traits are thus fixed. Note also that

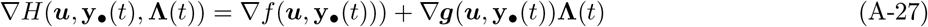

where

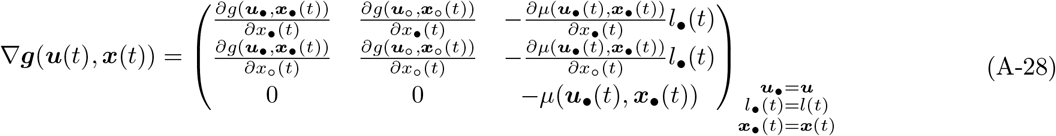

is a matrix, the gradient of vector ***g*** (hence *∇****g***(***u*, y**_*•*_(*t*))**Λ**(*t*) is the multiplication of a matrix by a vector). Finally,

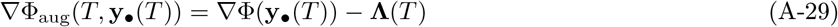

where

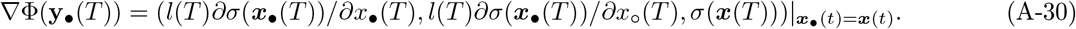

Now if we set

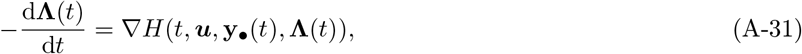

then the second term in the integral of eqs. (A-25)–(A-26) vanishes. Thus, on substituting eqs. (A-25)–(A-26) into the inclusive fitness effect

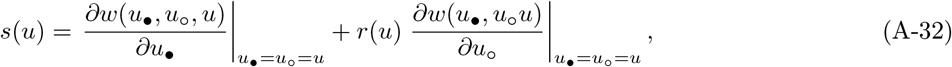

(eq. (2) of the main text) and simplifying shows that the inclusive fitness effect with discounted fitness (10) can be written as

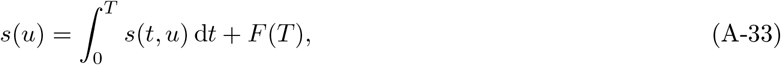

with

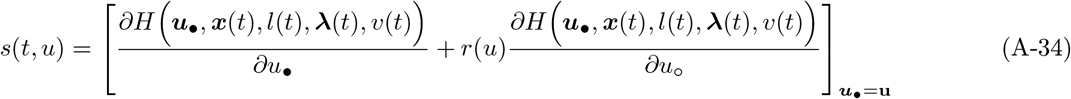

and

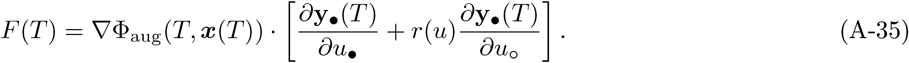

Since all state variables therein are evaluated at ***u***_*•*_ = ***u*** = (*u, u, u*), they satisfy the dynamics

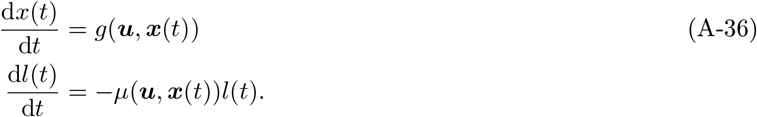

Exactly the same line of arguments can be applied to the integral appearing in the average fitness (12), with the difference that there is no state variable associated to survival and thus no costate. Carrying out the parallel calculations, then shows that the selection gradient (A-32) under average fitness can be represented as

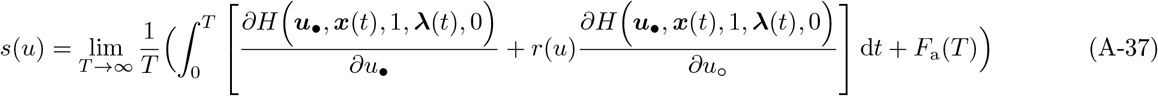

with

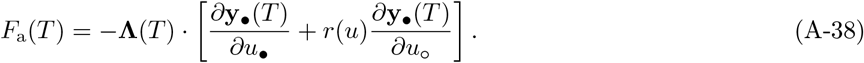

### Appendix B.2 The three expressions for the selection gradient

We here prove the three main text results using the generic represention for *s*(*u*) derived in Appendix B.1.

#### Appendix B.2.1 Result 1

We here prove result 1. Let us first consider the finite horizon case (*T < ∞*). Then, the boundary conditions for the costates are *v*(*t*) = *σ* (***x***(*T*)) and ***λ***(*T*) = *∇σ*(***x***_*•*_(*T*)) because this satisfies the definition of the costates as marginal changes of fitness at *T* (Box 1). These boundary conditions are also standard in control theory (Bryson and Ho, 1975; Dockner et al., 2000; Athans and Falb, 2007; Sydsaeter et al., 2008; Aseev and Kryazhimskiy, 2008; Weber, 2011) and imply that *∇***Φ**_aug_(*T*, ***x***(*T*)) = (0, 0, 0). Hence, it follow from eq. (A-35) that *F* (*T*) = 0, whereby the first line of eq. (A-33) reduces to eq. (17) and since together eq. (A-31) and eq. (A-36) are equivalent to eq. (14), this proves the finite horizon case of result 1.

Let us now consider the infinite horizon case (*T → ∞*). Owing to the assumptions of dominating discount, i.e., lim_*T→∞*_ ***λ***(*T*) = (0, 0) and that lim_*t→∞*_ *l*_*•*_(*t*) = 0, whereby lim_*t→∞*_ *∂l*_*•*_(*∞*)*/∂u*_*•*_ = lim_*t→∞*_ *∂l*_*•*_(*∞*)*/∂u*_*○*_ = 0, we have

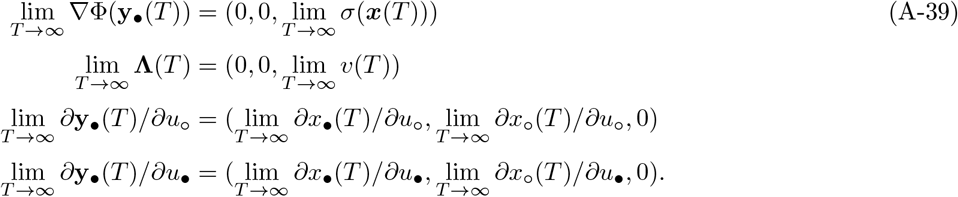

Substituting into eq. (A-35) shows that lim_*T→∞*_ *F* (*T*) = 0 (provided all entries remain bounded). Thereby eq. (A-33) reduces to eq. (17), which proves the infinite horizon part of result 1.

#### Appendix B.2.2 Result 2

We here prove result 2, which relies on the assumption that the initial condition of the state variable is given by the equilibrium 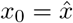 satisfying 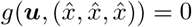 at ***u*** = (*u, u, u*). With this and owing to the fact that the selection coefficient *s*(*t, u*) in eq. (A-34) is evaluated at state ***x***(*t*) and thus at 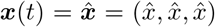 for all *t ∈* [0, *∞*), allows us to write eq. (A-34) as

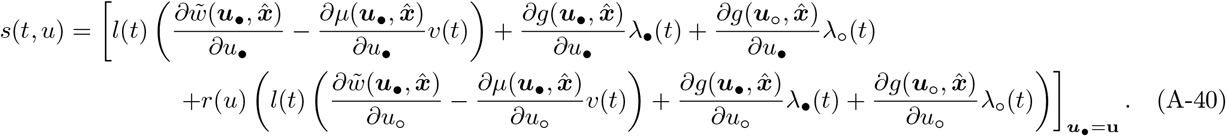

All vital rates and transition functions are here independent of time, since they are all evaluated at 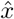. Hence, substituting eq. (A-40) into eq. (A-33), the vital rates and transition functions can be factored out from the integral and since an infinite time horizon with dominating discount is assumed, we have *F* (*T*) = 0. Therefore

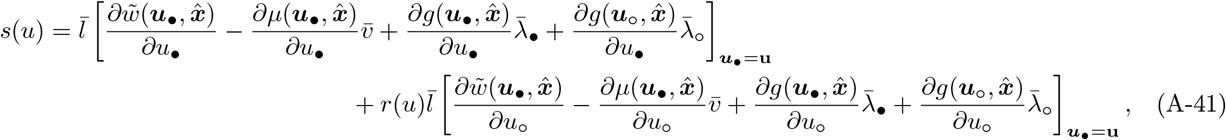

where 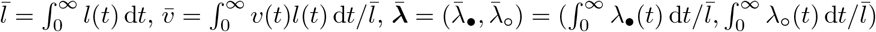. Using the definition of the Hamiltonian function, this can be equivalently written as

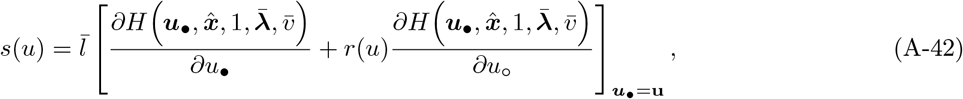

which gives eq. (21) for 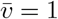.

To complete the proof of result 2, it thus remains to prove that 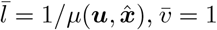, and 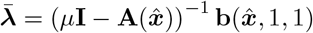.

For this, let us first note that under the assumptions underlying result 2, which involves dominating discounts, eq. (14) reduces to

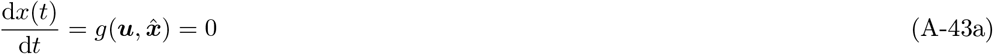

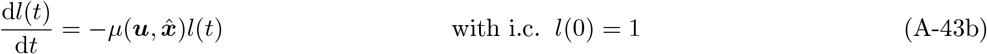

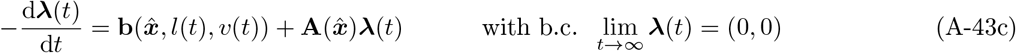

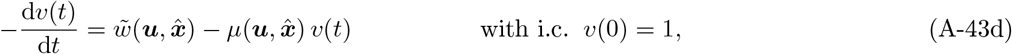

where we used the demographic consistency relation of Box 1 to obtain the boundary condition of the reproductive value. Straightfoward integration of eq. (A-43c), yields 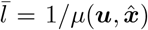. And since survival and effective fecundity is constant, we have 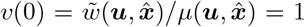. (Box 1), and so *v*(*t*) = 1 for all *t* satisfies eq. (A-43d), and thus 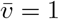. In order to prove the third equality, let us use the vectors

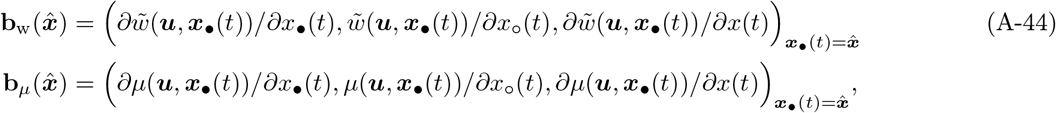

whereby eq. (15) can be written

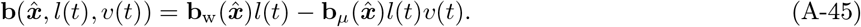

Substituting this into eq. (A-43) and intergrating yields

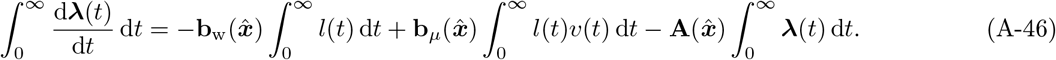

From the fundamental theorem of calculus 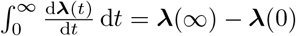 and so we can write

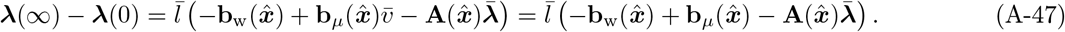

Since ***λ***(*∞*) = 0 by the assumption of dominating discounts, we need an expression for ***λ***(0) to obtain 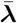. To than end, insert eq. (A-45) into eq. (A-6), which, for *t* = 0 and *ζ* = *T* with *T* = *∞* and ***λ***(*∞*) = 0, yields

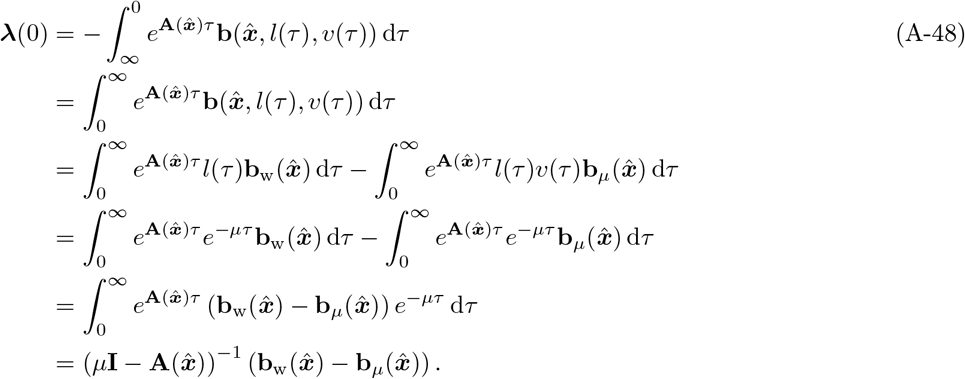

The second inequality follows from interchanging the boundaries, the third from inserting eq. (A-45), the fourth from inserting *l*(*τ*) = *e*^*−μτ*^ and *v*(*t*) = 1 (and using the shorthand notation 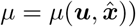, the fifth from reanranging, and the sixth from noting that the fifth line defines a Laplace transform and so for a constant vector **b** (Athans and Falb, 2007, p. 140). Inserting this into eq. (A-47) and solving for 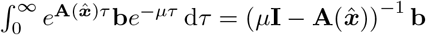 finally yields

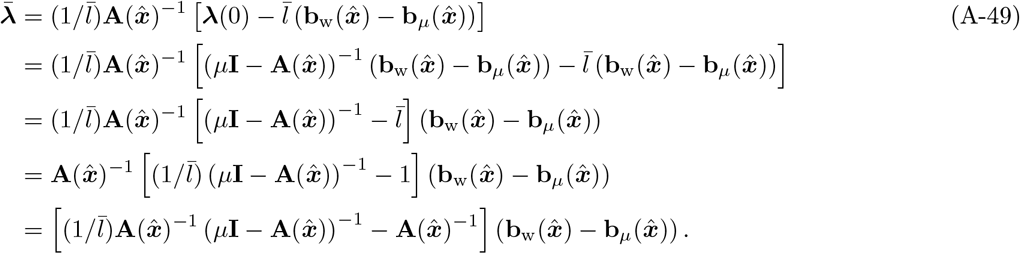

In order to further simplify this expression, note that 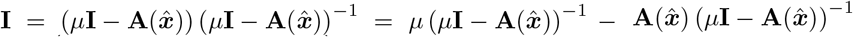. Hence, we have 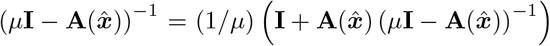 and on substituting into the last line of eq. (A-49) and using 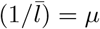 produces

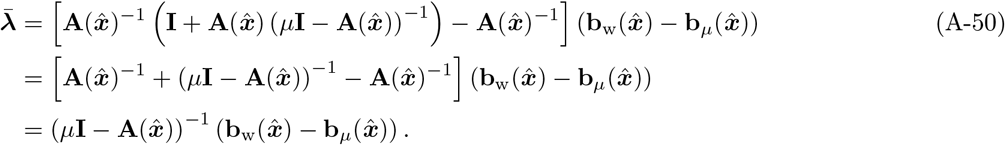

Using eq. (A-45) and reminding that 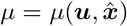, we finallly obtain 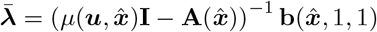, as needed to be proved.

#### Appendix B.2.3 Result 3

We here prove result 3, which assumes that given any initial conditions *x*(0) = *x*_0_ and ***λ***(0) = ***λ***_0_ at *t* = 0 for the states and costates, the variables converge to a unique equilibrium, 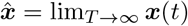, and 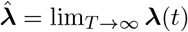, which is hyperbollically stable (*sensu* Hirsch et al., 2004). Then, owing to the facts that (a) *F*_a_(*T*) is finite in eq. (A-37) and (b) invoking the ergodic theorem, which informs us that the time average is the spatial average (Karlin and Taylor, 1975), then eq. (A-37) can be reduced to

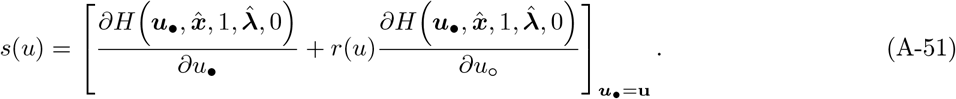

This proves the required result, since from eq. (14) the equilibrium 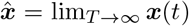 satisfies 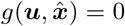 and the equilibrium 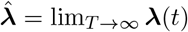 satisfies 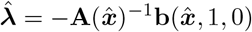.

## Notes

### Competing Interest Statement

The authors have declared no competing interest.

### Summary of Updates

The paper has been edited to improve the explanations and English. An new equation, namely eq.20 has been added and a new Appendix, namely Appendix A.2 has been added that explained in detail the assumptions of dominating discount.

